# Strain-level sample characterisation using long reads and MAPQ scores

**DOI:** 10.1101/2020.10.18.344739

**Authors:** Grace A. Hall, Terence P. Speed, Christopher J. Woodruff

## Abstract

A simple but effective method for strain-level characterisation of microbial samples using long read data is presented. The method, which relies on having a non-redundant database of reference genomes, differentiates between strains within species and determines their relative abundance. It provides markedly better strain differentiation than that reported for the latest long read tools. Good estimates of relative abundances of highly similar strains present at less than 1% are achievable with as little as 1Gb of reads. Host contamination can be removed without great loss of sample characterisation performance. The method is simple and highly flexible, allowing it to be used for various different purposes, and as an extension of other characterisation tools. A code body implementing the underlying method is freely available.

---

The ability to identify and quantify organisms within a sample underpins many fields of research, medicine, and agriculture. While cell culture and biochemical tests have historically been used to perform this type of sample characterisation, the greater speed, breadth, and resolution offered by DNA methods has made them highly attractive in recent years. Improvements to DNA sequencing technology and analysis software has driven new understanding of the microbiome's role in physical and mental health^1,2^, and has led to the creation of new microbiome-focused therapeutics^3,4^. DNA sequencing-based tests have been employed to diagnose patient infections, where they show higher accuracy and sensitivity than previous methods at a faster turnaround time^5^. While species-level characterisation has spurred progress in these fields, ambiguous strain-level results still hinder our ability to gain new insight into microbiome dynamics, and restrict the use of sequencing-based methods for pathogen identification. Strain-level sample characterisation – which strains are present, and their relative abundance -is now an area of active research, and may lead to a wave of new knowledge and opportunities once proper methods are discovered.

Marker-based approaches such as 16S rRNA characterisation were initially the most viable methods. An extensive range of organisms could be detected due to high availability of reference sequences and broadly applicable PCR primers, allowing these markers to be amplified, sequenced, then matched to a reference database. Marker based approaches target one or more genes, and use single nucleotide polymorphisms (SNPs) and small insertions / deletions (indels) to differentiate between taxonomic clades. While this approach can provide species level characterisation, inadequate genetic difference between strains in these marker regions has prevented strain-level resolution. Whole genome sequencing (WGS) approaches have recently provided better strain-level results as the entire genome of an organism can be used for differentiation. WGS methods have become more applicable to microbiome research and pathogen identification over time as the number of complete reference genomes publicly available has grown.

Approaches to sample characterization based on either short^6,7^ (including synthetic long read techniques^8,9^) or long read approaches are both effective. WGS based tools such as StrainPhlAn2^10^ for short reads, MetaMaps^11^ for long reads, and Wimp^12^, Kraken2^13^, and Centrifuge^14^ which may accept either length, all continue to expand our knowledge of the human microbiome. Due to the high per-base accuracy and market dominance of short read data, SNPs and small indels have historically been the focus for characterisation tools^12–14^ in the last decade. Both alignment-based approaches such as StrainPhlAn2, and heuristic approaches such as Kraken2 can be employed to probe this information. Despite a much higher per-base error rate (roughly 10% at current date), most long read characterisation methods tend to the same approach. Error-correction is frequently used in an attempt to compensate for the lower base-accuracy, where a common approach is to compare reads against one another to generate an error-corrected consensus sequence^15^.

The current challenge to strain level characterisation is the high level of genome-wide sequence homology between similar strains. Two different bacterial strains may differ by only a handful of SNPs and indels - a rate of less than 1 change per 100 kbp, hindering progress. Regardless of sequencing technology, most sample reads cannot be uniquely classified to a single strain, as there may be little if any sequence variation captured. This results in a swathe of similar organisms being reported alongside each true sample strain when the sample is characterised to strain level.

Aside from SNPs and small indels, long reads provide another avenue to exploit – the ability to easily identify structural variants (SVs). Long reads provide a much more powerful tool for recognising and characterising structural variants (SVs) - elements longer than 50 or so bases - than the reads from NGS technologies^16–18^. Tools that allow the extraction of a range of different structural variants from long reads are available^19^, and their use in human health studies^20, 21^ have been recently documented. The importance of SVs in phylogeny^22^, and therefore in the differentiation of organisms, is well-established, and exploitation of long read technology in this field is rapidly progressing^23^.

We take a new approach to sample characterisation. Rather than focusing on how to remedy the deficiencies of long reads, we directly exploit their strengths - namely that they are long, even if error prone, and their ability to sample large genomic regions provides ready access to structural information. Reads spanning discriminating structural features provide clear differentiation between strains, and are used to identify the true sample strains over other, highly similar organisms. The method has been implemented in our code body, NanoMAP (**Nano**pore **MAP**Q characterisation tool).

## RESULTS

### MAPQ Scores Can Differentiate Highly Similar Strains

Microbial genomes are highly dynamic. Entire sections of DNA are often inserted, deleted, inverted, copied, or otherwise moved within and between microorganisms. While SV types such as insertions and deletions have been indirectly used by characterisation tools through identifying clade-specific DNA^24^, structural variation can also create clade-specific arrangements of DNA. Even in the case of cut-paste variants where the DNA content is not modified, the sequential arrangement of DNA is altered. These genetic rearrangements can generate discriminating regions of DNA where only a single strain possesses a particular structural arrangement. Reads sampling these unique regions align markedly better to the organism’s true reference genome over those belonging to other strains. Mapping quality (MAPQ) scores are used to detect these reads, and their use underpins our method for sample characterisation.

MAPQ scores provide a measure of the confidence that a read actually originates from the position it is aligned to. Reads which align well to only a single location in the reference sequence have an alignment with high MAPQ score, while reads which have multiple locations of equal best alignment have MAPQ score equal to zero for all alignments of the read. They were initially designed to indicate the chance a read had been misplaced in a single organism’s reference genome, but if we treat a whole database of strains as a single reference genome, a high MAPQ score alignment can signal a read which has been mapped uniquely to a single genome in that database. This scenario can be established by simply concatenating the genomes of interest (the database) into a single reference metagenome. Reads reporting a high MAPQ score alignment (HMQ reads) have a single best mapping location in the metagenome, implying they align markedly better to one strain over all others. Figure 1a demonstrates this process, showing how a single SV can differentiate between strains. Figure 1b provides our implementation of this method for sample characterisation, and Figure 1c provides an indicative set of discriminating data generated by application of this method using NanoMap.

**Figure 1.**
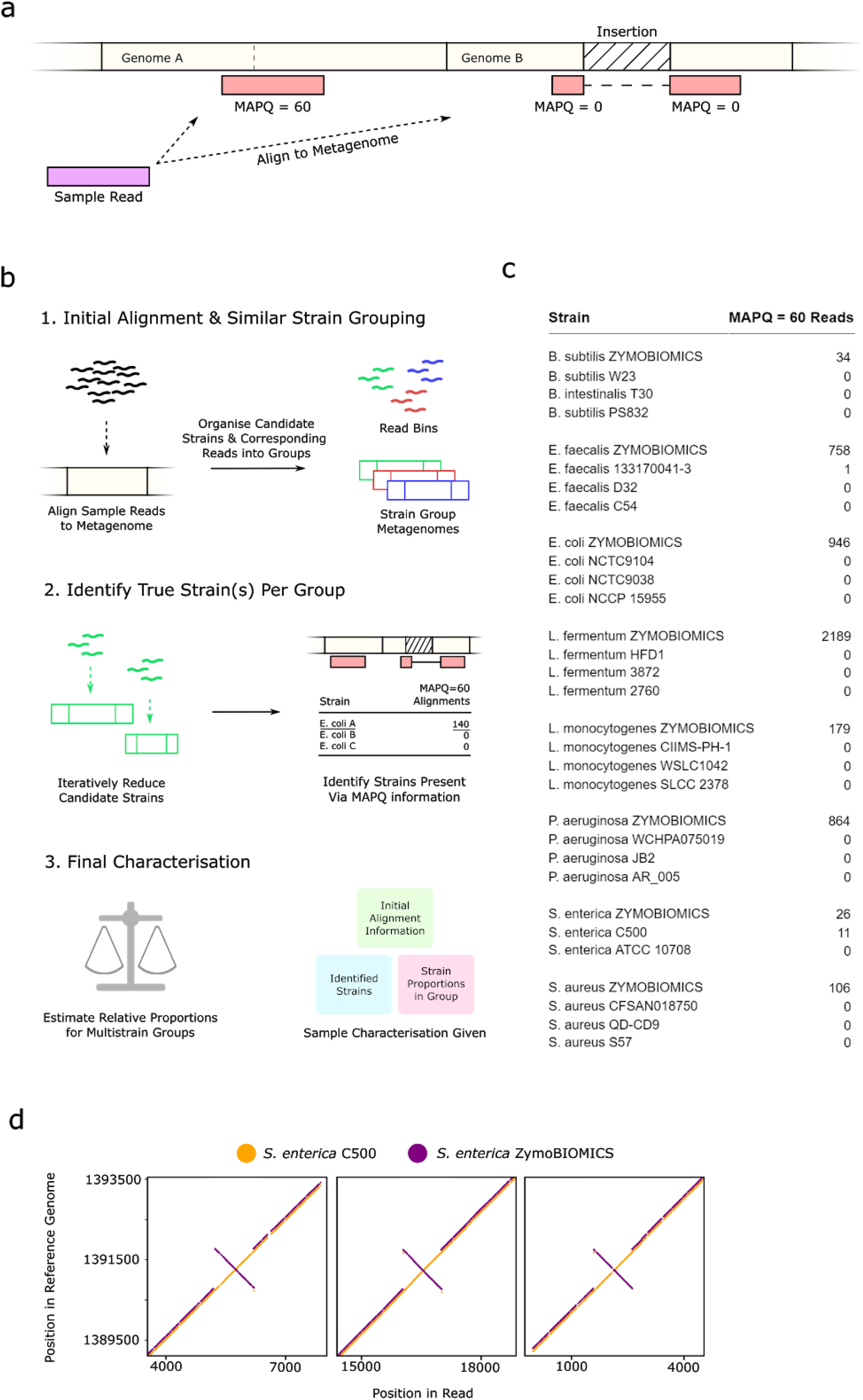
Summary of the key elements of our approach, and data from characterisation of the ZymoBIOMICS Microbial Community Standard product. **a)** Alignments of a sample read (originating from organism with reference genome A) to a metagenome. Alignment to the correct reference genome appears full length, while alignment to a similar strain’s reference genome produces a split alignment. The alignment to genome A may possess a high MAPQ score due to poor genome B alignment. **b)** Overall program workflow that has been implemented in the NanoMAP code. **c)** Number of MAPQ=60 read alignments per strain during characterisation of theZymoBIOMICS Microbial Community Standard product. The top 4 strains (by MAPQ=60 alignment count) in the final round of alignment from each candidate strain group are shown. **d)** Kmer dotplots for 3 of the 11 off-target (S. enterica C500) HMQ reads vs the ZymoBIOMICS and C500 strains. The C500 reference genome was shifted 1615000 bp to place it in alignment with the ZymoBIOMICS reference genome. A recent inversion in the ZymoBIOMICS organism is not reflected in its supplied reference genome.

This method is simple in approach and general. Any read alignment tool which reports MAPQ scores can be used, with our aligner of choice being minimap2^26^. While the majority of discriminating reads seem to be produced by the presence of structural variation, any form of genetic uniqueness can contribute. SNPs and indels can generate HMQ reads if enough genetic difference of these types is available in a given region, which may be relevant for some taxonomic clades. No set database is required, as only sample reads and reference genomes are needed for characterisation.

### Single SVs Can Provide Strain Level Discrimination

The use of MAPQ scores provides powerful strain discrimination. Figure 1c displays the final set of information used by NanoMAP during runtime to identify sample strains. A clear distinction between correct and incorrect strains is seen in the HMQ read count for ZymoBIOMICS strains and other, highly similar organisms. Strains are listed by HMQ read count, demonstrating that ZymoBIOMICS strains often have hundreds if not thousands of HMQ reads while other strains have zero. Aside from *S. enterica,* all expected strains are clearly distinguishable as the true sample strain for each species.

After investigation into the off-target *S. enterica* C500 HMQ reads, it was discovered that the reference genome for the correct *S. enterica* strain is out of date. The *S. enterica* ZymoBIOMICS strain appears to have recently experienced an inversion in a phase variation control region called *hin*^27^. As a result, the supplied ZymoBIOMICS reference genome for this strain does not reflect the change. Phase variation permits fast adaptation to a changing environment via switching the expression of certain genes on or off, and in this case the orientation of *hin* governs the expression of flagellar proteins. The reference genome for strain C500 has *hin* in an orientation which matches that of the *S. enterica* ZymoBIOMICS cells present in the sample, causing the 11 off-target HMQ reads. Figure 1d shows how these reads would align to both these strains, showing kmer matches between the read and *S. enterica* strain C500 in yellow, with *S. enterica* ZymoBIOMICS in purple. All off-target HMQ reads covered the inverted *hin* region.

MAPQ scores can differentiate between strains even in the case of a single SV being the only genetic difference. Aside from the single inversion which caused off-target HMQ reads explored above, this principle is clearly demonstrated in Figure 2a. HMQ read count data produced during characterisation of an ATCC mock microbiome shows sample strains *H. pylori* 26695 and *E. faecalis* V583 being distinguished from other organisms. The strains listed below *H. pylori* 26695 and *E. faecalis* V583 in the table are derivatives of these true sample strains, and differ by as little as 1 SV to the original organism. The transformations used to create the derivative strains are shown in the comments column. A single gene replacement between *H. pylori* 26695 and *H. pylori* 26695-dR is capable of uniquely identifying 26695 as the true sample strain. Similarly for *E. faecalis*, a single 280 bp insertion in strain VE18379 provides enough discrimination to identify V583 as the true sample organism.

**Figure 2.**
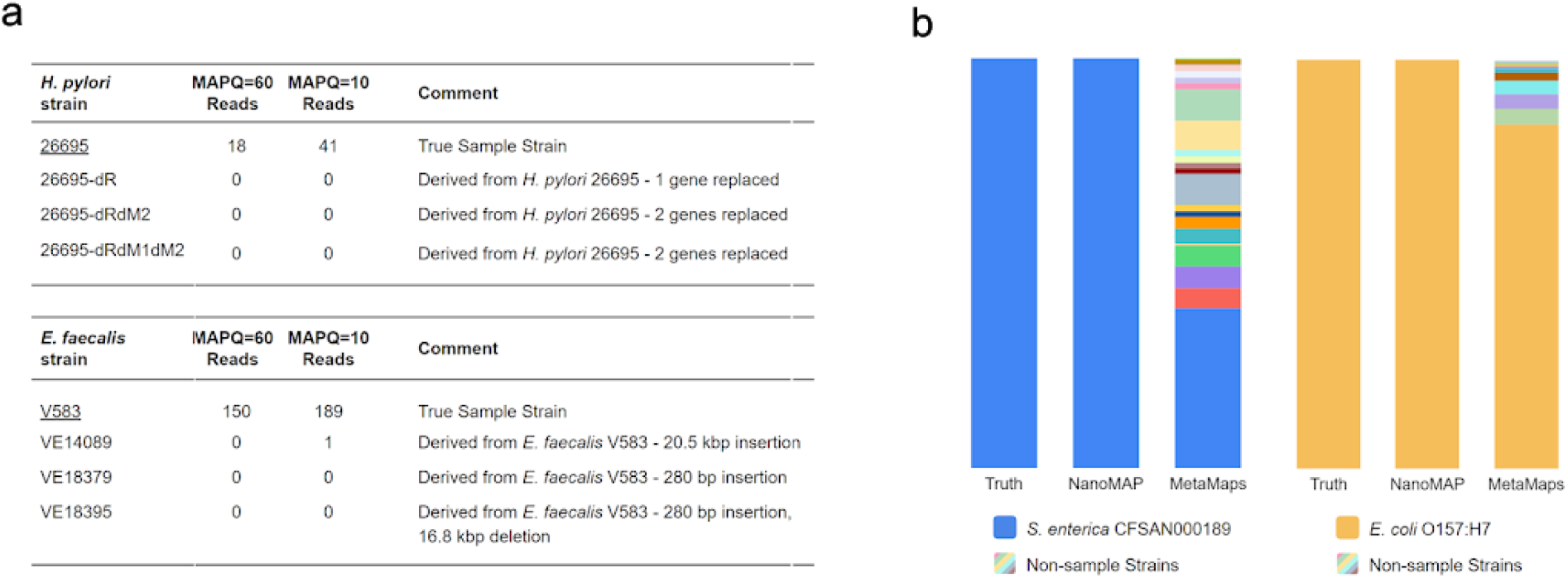
Accurate strain-level characterisation of multiple organisms. **a)** Table of final round HMQ read counts for two strains present in the ATCC mock microbiome sample. The true sample strains, H. pylori 26695 and E. faecalis V583, differ from other possible strains by at most 2 SVs **b)** NanoMAP and MetaMaps characterisation of two samples, each sample containing a single organism. NanoMAP reports only the true strain in both cases, while MetaMaps reports a large number of incorrect strains alongside the sample strain.

Figure 2b demonstrates true strain-level characterisation for two single-strain samples using this method. Two samples, one containing a single *S. enterica* strain and the other containing a single *E. coli* strain were characterised by NanoMAP and MetaMaps using a consistent database for both programs. For the *S. enterica* sample, the database consisted of all complete *S. enterica* assemblies deposited on RefSeq (as of 28/09/2020). For the *E. coli* sample, the database consisted of all complete bacterial (19,077) and fungal (12) genome assemblies on RefSeq at the same date, with the addition of human reference genome hg38 (hence known as the RefSeq b+f+h database). The reported composition returned by each tool is shown as a stacked bar chart. NanoMAP reported only the correct strain for each sample providing accurate characterisation, while MetaMaps reported an ambiguous characterisation consisting of multiple strains for each sample.

### Iterative Alignment Provides Access to Strain Discriminating Data

The following two sections provide a brief overview of our code body, NanoMAP, which implements this method. Fuller details can be found in the Methods section.

NanoMAP uses multiple rounds of alignment in its approach. Alignment methods for sample characterisation have been criticised as time-consuming when a large reference database is used. To overcome this issue a multi-step method is taken, where a rough characterisation is initially sought, followed by a more targeted approach. An initial alignment at *k*mer-level, rather than base-level accuracy, maps all sample reads to the full reference metagenome. This allows a shortlist of candidate strains (strain group) for each real sample strain to be derived using the approximate mapping information. Multiple rounds of base-level accuracy alignment are performed for each strain group to narrow the set of candidate strains and provide necessary MAPQ information. The use of approximate mapping in the initial alignment step drastically reduces runtime, with no current evidence of degraded performance.

The role of initial full database alignment is twofold - to confine each sample organism to a small set of possible strains, and to bin reads by the organism from which they appear to originate. The alignment output file is processed to provide this information. Reads which map poorly to the full metagenome are removed through minimum percent identity and alignment length cutoffs. Short reads (< 1000 bp) are also discarded during this process. Each read is then classified to a subset of database organisms by comparing all alignments of a given read. The rate of strain co-classification is recorded during this stage, and is used to identify which strains should appear together as candidate strain groups. This approach is flexible in terms of taxonomic labels as strain groups are allowed to contain strains across species designations, and multiple strain groups can be formed within a single species if the sample contains multiple organisms of the species.

Reads are binned by the most appropriate strain group according to their classification. Any read which was classified to reference genomes belonging in two separate groups is discarded. The naive DNA abundance (%) of each group is calculated as the total number of base pairs in reads binned to the group, divided by the total number of base pairs in all read bins. To prepare for secondary alignment, a fastq file is generated from each read bin, and a targeted reference metagenome is built for each candidate strain group.

Iterative alignment of each read bin to its corresponding metagenome allows the true sample strain(s) to be identified. For a given strain group, the list of candidate strains is narrowed each iteration until the correct organisms(s) can be identified via HMQ read counts. An iteration consists of three steps: Base-level alignment of the read bin to the strain group’s current metagenome, output file processing, and halving the remaining candidate strain pool by removing unlikely candidates. A new metagenome is constructed from the genomes of the remaining strains. If a single strain is clearly evident in the group, or ≤ 4 strains remain, the halving process is skipped, and strain identification is performed using HMQ read count information. Hard cutoffs, and ratios of HMQ counts are used to conclude which of the final strains are truly present in the sample. Often a single sample organism is present in each strain group, but multiple sample organisms can appear together if they possess high sequence homology. In these cases, the relative abundance of each identified strain needs to be estimated.

### Proportion Estimation for Strains in a Strain Group

When NanoMAP identifies multiple strains within a group, their relative proportions must be estimated. As these strains were placed in the same group, they will possess a high level of sequence homology. This poses a challenge to abundance estimation, as most reads binned to the strain group will align (and therefore be classified) equally well to each identified organism’s reference genome. If we estimate abundance naively as above, all identified strains will display roughly equal sample proportion due to these equal-best alignments, even when the true proportions within the group differ greatly. Accurate strain-level characterisation under these circumstances requires more than a simplistic approach.

To overcome this issue, we use a probability model to estimate classification frequencies. Genomic reads of identified strains are simulated, aligned, then classified to these strains, allowing a model to be generated to estimate read classification probabilities. For a given strain group with multiple identified strains, we take the observed classification frequencies of the strain group’s read bin, and use the probability model generated to estimate the proportion of reads from each identified strain within the group.

To illustrate, an example where 3 strains (A, B and C) have been identified will be used. Our read set is the fastq file of *n* reads for the group being assessed, and the possible classifications for each read are {A, B, C, AB, AC, BC, ABC}. A read is classified as A if its best alignment among the three genomes is unique to genome A, while it is classified AB if its best alignment among the three is an equally good alignment to genomes A and B, but not to genome C. Similarly for AC and BC, while a read is classified ABC if its best alignment is an equally good alignment to all three genomes A, B and C. Now replace the strain labels A, B and C by *i*=*1,2* and *3*. Each read receives one of the seven classification*s* which we now label *j* = 1,…,7. Our observed data are then the counts *{n_j_},* where *n*_*j*_ is the number of reads receiving classification *j*. The probability model underlying our estimation procedure supposes that each read has an unknown probability *p*_*i*_ of being from strain *i,* and, given that a read is from strain *i*, it has probability *t*_*ij*_ of receiving classification *j*. The probabilities *{t_ij_}* are estimated by simulation. With the combination of the estimated probability matrix *{t_ij_}* and the observed classification counts *{n_j_}* for the read set, an EM-algorithm for estimating the probabilities *{p_i_}* can be derived, and that is what we use. See Methods for further details.

### Unambiguous Strain-Level Characterization of Two Microbiome Products

To test the viability of this method for more complex samples, two mock microbiome products were characterised. The ZymoBIOMICS Microbial Community Standard, and ATCC MSA 2006 (henceforth referred to as ZYMO and ATCC samples respectively) products have known composition, and were characterised to strain level by NanoMAP and MetaMaps. For both mock microbiomes, Nanopore sequence data was already available. The read sets of these products were downsampled to 1Gb of sequence data, and this subset was used for characterisation. The aforementioned RefSeq b+f+h database was used for both NanoMAP and MetaMaps characterisation, with the addition of ZymoBIOMICS and ATCC reference genomes for sample strains. Any duplicate reference genomes of sample organisms were manually removed from the database after these additions (Supplementary Information).

Figure 3 shows NanoMAP providing accurate and clear characterisations for both mock microbiomes. For the ATCC mock microbiome, NanoMAP reported no non-sample strains, while MetaMaps reported 16 at sample DNA abundance > 0.1% (Figure 3c). These 16 extra strains represent 8.2% of sample composition in the MetaMaps characterisation. MetaMaps provided good characterisation for most species, but struggled with *H. pylori* and *S. enterica*. These species were challenging for MetaMaps as organisms which are highly similar to the sample strain are present in the database. Sample composition reported by each tool is seen in Figure 3a, and shows high concordance between NanoMAP and MetaMaps despite differing from the theoretical composition.

**Figure 3.**
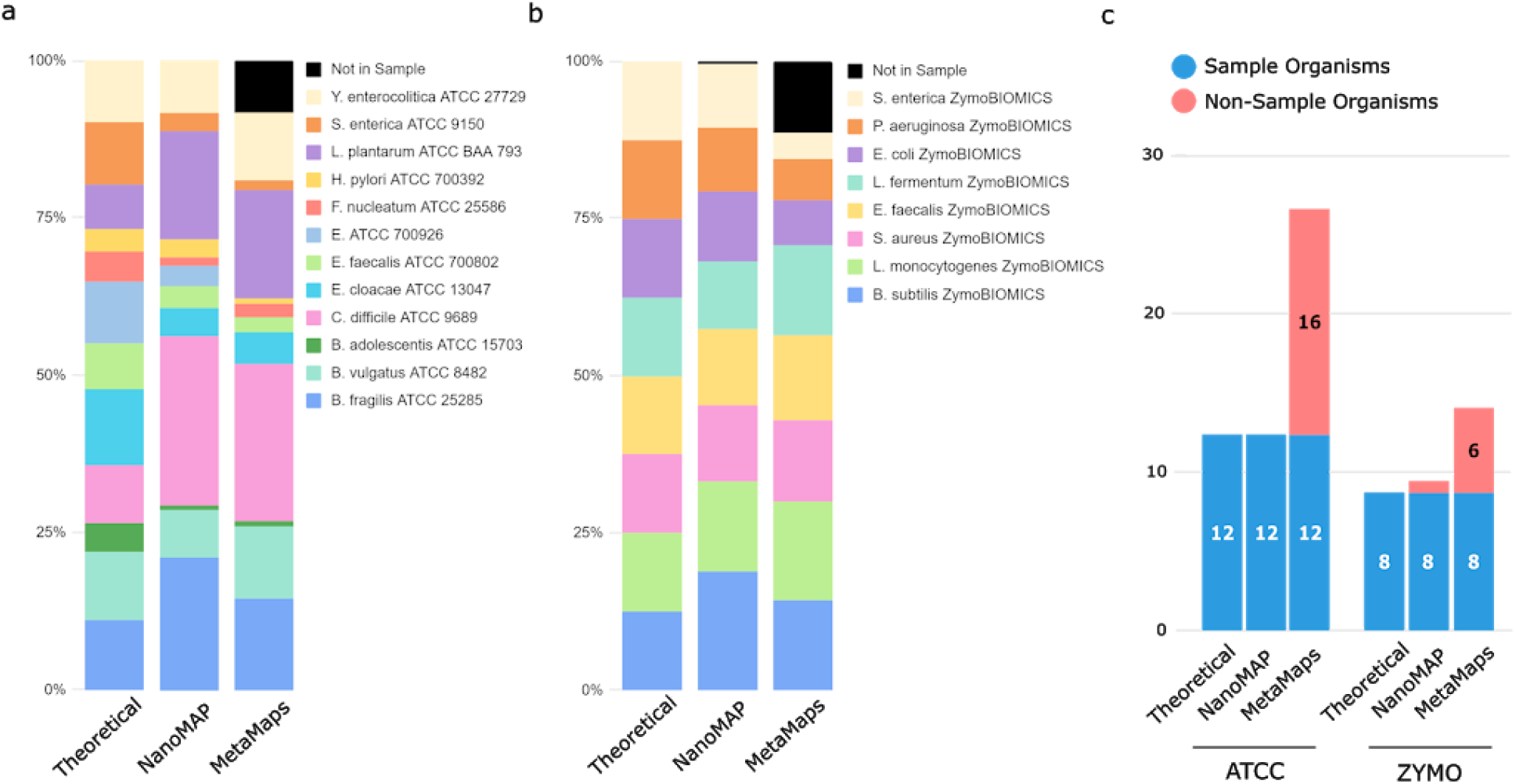
NanoMAP and MetaMaps characterisations of ATCC and ZYMO samples. **a)** The theoretical composition of ATCC shown with sample composition reported by NanoMAP and MetaMaps. **b)** Theoretical composition of the ZYMO sample shown with those reported by NanoMAP and MetaMaps. **c)** Number of sample and non-sample strains reported by NanoMAP and MetaMaps for the ZYMO and ATCC samples. Number of sample strains identified is shown in blue, with the number of non-sample strains reported shown in red.

For the ZYMO sample, NanoMAP reported one non-sample strain, *S. enterica* C500, as present, while MetaMaps reported 6 (Figure 3c). The composition for the ZYMO sample, seen in Figure 3b closely matches the theoretical values. Multiple other independent characterisations of this mock microbiome^30,31^ mirror the inflated abundance for *B. subtilis*. Many explanations of this are possible, including that the real sample composition diverges from theoretical, or that the sample preparation methods cause this effect. Nevertheless the compositions reported by NanoMAP and MetaMaps appear to be reasonably accurate. For both samples, MetaMaps strain-level composition is slightly altered from NanoMAP. This is mostly due to MetaMaps being unable to resolve some sample organisms to the single correct strain. In these cases the non-sample strain(s) contribute to the ‘Not in sample’ segment seen in black.

The genome database must contain a single copy of each sample organism’s reference genome. As strain-level characterisation relies on little genetic diversity to differentiate strains, these reference genomes must be current and reflect the true genome of sample organisms. This is true for all characterisation tools, but especially so for NanoMAP. Table 1 shows the different outcomes that may be seen depending on different database states, and highlights the need for a good-quality, non-redundant collection of reference genomes. It also demonstrates that different types of deficiency in the database with respect to a strain group of interest can be diagnosed. *L. plantarum* ATCC BAA-793 reads, acting as a sample, were aligned to a metagenome consisting of 6 complete *L. plantarum* genome assemblies. The correct strain is identifiable in the ‘Single Copy’ scenario, but not the ‘Redundant’ or ‘Not Present’ situations. NanoMAP requires the database to be consistent with the ‘Single Copy’ scenario, where a single copy of each sample organism’s reference genome is present. The other scenarios demonstrate that strain-level characterisation is not possible using MAPQ score information in the case of a database redundancy, or a missing reference genome.

**Table 1.** Indicative HMQ read counts for three database scenarios using a sample of L. plantarum ATCC BAA-793 reads. The middle ‘Single Copy’ scenario denotes a situation where the database contains a single copy of the correct reference genome for the sample reads. ‘Redundant Copy’ shows how a redundant copy of the correct reference genome leads to zero HMQ read alignments to database genomes. When the correct reference genome does not exist in the database, as shown in the ‘Not Present’ table, many non-sample strains exhibit HMQ alignments due to local regions of sequence homology to the correct reference genome.

| Redundant |  | Single Copy |  | Not Present |  |
| --- | --- | --- | --- | --- | --- |
| Strain | MAPQ=60<br>Reads | Strain | MAPQ=60<br>Reads | Strain | MAPQ=60<br>Reads |
| *L. plantarum* ATCC BAA-793 | 0 | *L. plantarum* ATCC BAA-793 | 4469 | *L. plantarum* strain HEAL19 | 773 |
| *L. plantarum* ATCC BAA-793 Copy | 0 | *L. plantarum* strain HEAL19 | 0 | *L. plantarum* strain BLS41 | 495 |
| *L. plantarum* strain HEAL19 | 0 | *L. plantarum* strain BLS41 | 0 | *L. plantarum* strain LM1004 | 427 |
| *L. plantarum* strain BLS41 | 0 | *L. plantarum* strain LM1004 | 0 | *L. plantarum* strain DSR_M2 | 362 |

### MAPQ Method Shows Wide Applicability

Given an appropriate database, this method appears to allow strain differentiation in most situations. To test its applicability to other organisms and taxonomic clades aside from those encountered during development, we investigated whether each bacterium with a complete genome assembly available on RefSeq (19,077) could be distinguished from any other bacterium in this set via HMQ read count. An all-vs-all approach was not pursued in this situation due to the large number of organisms in this group. Rather, we made the assumption that any bacteria which can be distinguished from its most closely related organism via MAPQ information should also be distinguishable from less related organisms using the same method. For the following, a single organism is used as an example, with the same procedure being carried out for all 19,077 complete bacterial genomes in the RefSeq bacteria group.

For a given query genome, its most similar counterpart was determined through alignment percent identity. 1000 genome fragments of length 3000bp were first extracted from the query with uniform coverage and spacing across its length. These fragments were then aligned to a database containing all RefSeq complete bacterial genomes to base-level accuracy using minimap2. All alignments for the query fragments were grouped by reference genome to which they were being aligned. The top 20 reference genomes by number of alignments were retained and other reference genomes discarded. Query fragment alignments to the query reference genome itself were removed. For each reference genome exhibiting alignments, the median pid across its alignments was calculated, and reference genomes were ranked by this statistic. Reference genomes exhibiting median alignment pid > 99.999% were removed, as this SNP rate (approximately 1 in 100,000bp) is within the expected error for modern assemblies, and a proportion of these were assumed to be redundant copies of the query genome. After this step, the reference genome with the highest median pid was reported as the most similar counterpart to the query genome being assessed.

By this stage, each bacterium in our RefSeq database had been linked to its most similar counterpart. The ability for each organism to produce MAPQ=60 alignments to its own reference genome over the reference genome of its counterpart was then measured. For each query bacterium, a metagenome was constructed from its reference genome and the genome of its counterpart. 300 reads of the query bacterium were simulated with NanoSim^29^, using an error model trained on *S. enterica* ZymoBIOMICS read alignments to the corresponding *S. enterica* ZymoBIOMICS reference genome. The simulated reads were then aligned to this two organism metagenome. Reads with length < 1000bp, were removed, then the number of alignments with high MAPQ score to either organism was noted.

Figure 4a suggests that the MAPQ method may be able to discern true sample strains in the vast majority of cases. 99.75% of query / counterpart bacterium pairs produced at least one MAPQ=60 read alignment to the query genome over its most similar in-database counterpart from the pool of 300 simulated reads, with that proportion reaching 99.99% if MAPQ=10 alignments are also considered. For more marked differentiation, 98.49% of query bacterium displayed at least 10 MAPQ=60 alignments, with 99.98% displaying at least 10 MAPQ=10 read alignments. As expected, no HMQ reads to the counterpart rather than the query were witnessed across all comparisons, but this may not be the case if read errors such as chimeras, random reads and junk reads are introduced by the sequencer. This data suggests the use of high MAPQ score reads may be reliable for the majority of possible sample compositions.

**Figure 4.**
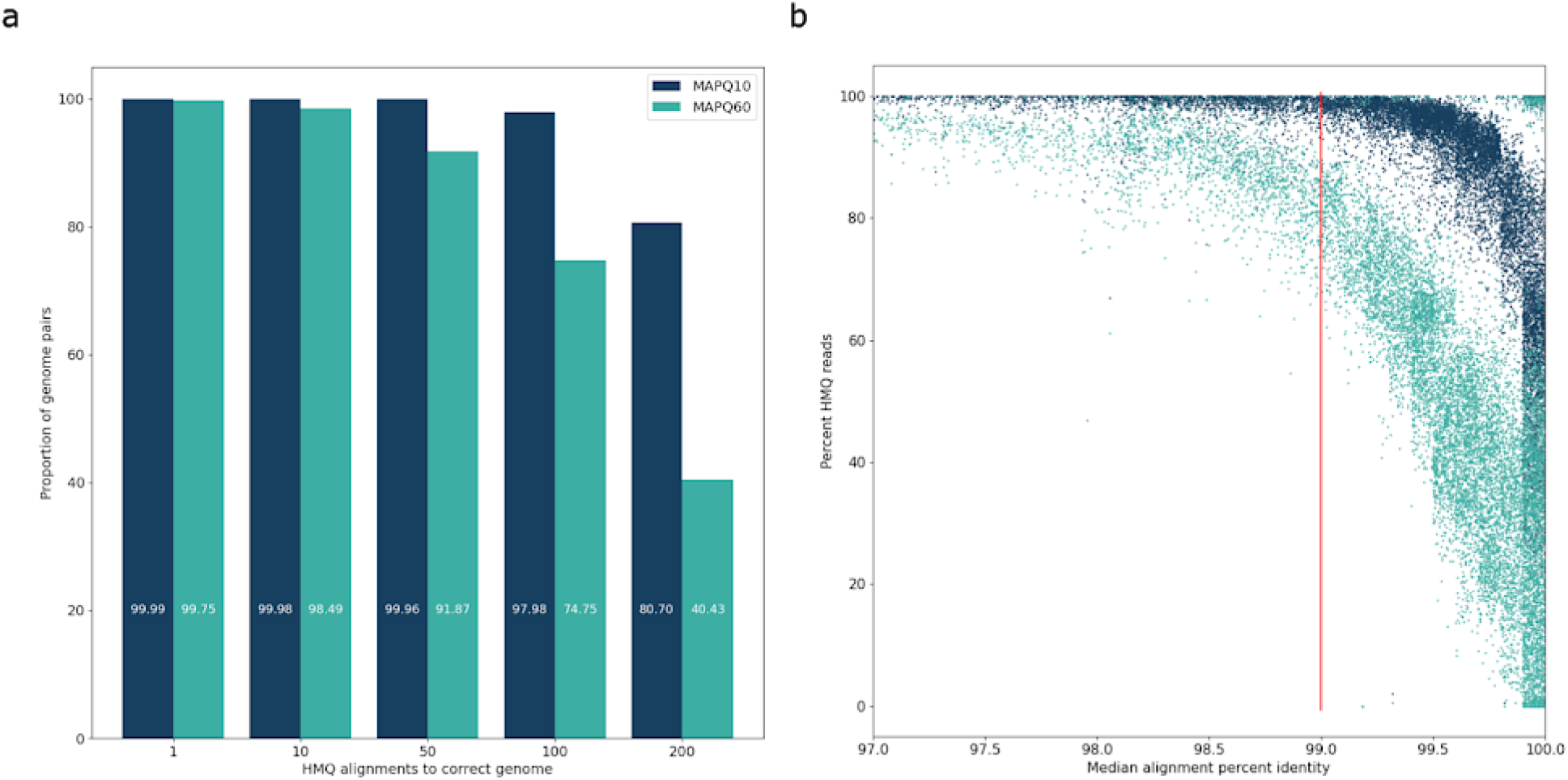
RefSeq database MAPQ analysis. Each query genome in the RefSeq bacterial database was paired with its most similar counterpart, as measured by median alignment pid between the query and counterpart organism. **a.** Proportion of query/counterpart pairs with equal or greater than **x** high MAPQ alignments to the query, from a set of 300 simulated reads. No HMQ read was witnessed to any counterpart genome in all comparisons. **b.** Sequence identity vs proportion of reads which had HMQ alignment to query. A high proportion of simulated query reads possess a high MAPQ score alignment to the query over the counterpart genome, even when the two genomes have greater than 99% sequence identity.

Figure 4b indirectly shows the relationship between sequence homology and MAPQ read generation. For query / counterpart bacterium pairs whose genomes diverge by roughly 1 in 200 bp, more than half of all simulated reads produced a MAPQ=60 alignment to the query (correct) genome and can be uniquely mapped. When considering a MAPQ score of 10, query / counterpart pairs approaching 1 base alteration per 1000 bp still display a large proportion of reads with MAPQ=10 alignment to the query. Above this level of sequence homology, most reads are multimapping rather than uniquely classified to the query, although the data in 4a shows that some distinguishing reads are usually present. The high proportion of reads with MAPQ=60 alignment despite high sequence identity between the query and counterpart further supports structural variation as the main generator of HMQ reads. As a MAPQ score of 60 for a read alignment theoretically represents a 1 in 1,000,000 probability of incorrect mapping location for the read, it is unlikely that a small number of distinguishing SNPs and indels which may have been sampled by a read cause such high confidence in mapping location. The 300 simulated query reads equate to 2.5 Mb of sequence data (0.25% of 1Gb read set), suggesting that strain-level identification may be possible down to very low sample abundances for some clades using this method.

### Sensitivity

The data in Figure 1c suggest this method can provide strain-level discrimination even for lowly abundance sample organisms. During ZYMO sample characterisation, the ZymoBIOMICS strains (each present at 12% sample abundance) generally had hundreds of MAPQ=60 reads, while other organisms of these species possessed zero. Assuming that the MAPQ=60 read count is proportional to the total number of reads sequenced for these strains, most ZymoBIOMICS strains would still have tens of MAPQ=60 reads if their sample abundance was reduced to less than 1%, permitting correct identification. Sensitivity fluctuates depending on organism, and Figure 1c shows that strain-level identification would only be possible to approximately 1% sample DNA abundance for the *B. subtilis* and *S. enterica* ZymoBIOMICS organisms.

A simple modelling approach provides further assessment of the sensitivity of this approach. We require at least one HMQ to declare one genome different from another. Then, following an approach similar to Lander and Waterman^30^, we can estimate the probability that a random sample of reads includes at least one read that crosses a structural variant boundary. This can be estimated by considering cases in which there are *n* structural variants between the pair of genomes, and requiring there to be a minimum amount of each read lying in both the structural variant region and the common region of the pairs of genomes that a sufficiently high MAPQ value is achieved. Such an analysis (Supplementary Information, Figure 1) shows that, with only one discriminating SV to the most similar other organism, and only 1Gb of sequence data, it is expected that - even at a relative abundance of 0.25% - a strain would be identified nearly 50% of the time. If there are 5 discriminating SVs this rises to strain identification more than 95% of the time. Thus the empirical results are supported by a more general, though simple, theoretical analysis.

## Discussion

The method presented here offers fine-grained identification of the microbial organisms present in a sample. Accurate characterisation of both single- and multi-organism samples may allow the method to be employed for a wide range of purposes including microbiome research, pathogen identification, and environmental sample analysis. While these possibilities are exciting, strain-level characterisation is only possible for strains present in the reference database. An increasing number of relevant genomes are being added to publicly available catalogues, providing more likelihood of encompassing the organisms present in a sample.

Researchers must be confident in the output provided by characterisation tools, as incorrect characterisation may divert research away from the best direction. An ideal tool would need to identify all strains with noteworthy abundance in the sample while reporting no incorrect organisms. While the method presented here still has much room for development, NanoMAP demonstrated these abilities for the ATCC mock microbiome sample, and was only prevented from doing so with the ZYMO sample due to an out of date reference genome.

The method’s ability to distinguish strains based on structural variation makes it useful for pathogen identification. Pathogenic and benign strains of a given species may only differ by a small number of relative SVs, requiring diagnostic tools to be able to discriminate using these features. NanoMAP use has demonstrated the ability to differentiate between strains when a single 280 bp SV was the only marked genetic difference. Host contamination needs to be removed for patient samples but poses no significant issue, as human reads are binned to a strain group consisting of the human genome during NanoMAP runtime and can be ignored.

This method can be extended to provide other functionality. If desired, NanoMAP can act as a final pass following another tool’s characterisation. The strains reported by the initial tool can be fed to NanoMAP, where it may be able to provide further strain-level information. In future, the use of MAPQ scores may be expanded to perform even greater tasks, such as providing the ability to monitor the evolution of microbiome organisms as they acquire structural variation, and to measure the impact of these genetic changes on human health. Database redundancies may also be identified by aligning fragments of database genomes back to the database itself, and viewing the resultant MAPQ score information. Table 1 demonstrates the expected MAPQ score information given the database contains a single copy, or redundant copies of a given reference genome, which could be used as a basis for this task.

Currently, the NanoMAP implementation of these ideas differentiates between strains via HMQ read counts. Rather than using these key reads to simply discriminate between strains, the genetic differences generating HMQ reads could be reconstructed. This may provide greater discriminating power, and error-correction potential. Off-target HMQ reads arising from reference genome errors, as seen in the Salmonella enterica data of Figure 1c with the associated inversion seen in Figure 1d, or those caused by read errors, such as chimeras, may be avoided using this approach.

## Methods

### Sample Characterisations

Four datasets were used in assessing the performance of NanoMap. The choice of datasets was guided by recent published work^11,13^ and to cover the different use-cases for this method. For the characterisation of each dataset, NanoMAP and MetaMaps were supplied an identical set of reference genomes to use as a database.

#### Two Mock Microbial Communities

The ZymoBIOMICS Microbial Community Standard Cat #D6300 and ATCC MSA-2006 are established datasets for assessing performance of microbiome analysis methods and tools. Nanopore sequence data is publicly available for both samples. The ZYMO read set, sequenced on the nanopore MinION device by Nicholls et al in 2019, was downsampled to 1 Gb for characterisation. Downsampling was performed by selecting a contiguous 1 Gigabase read chunk from part-way through the original fastq file. The ATCC read set generated by Moss, Maghini and Bhatt^31^ was downsampled in a similar manner. The database used for both samples was RefSeq b+f+h with the addition of ZymoBIOMICS and ATCC reference genomes for sample strains. Redundant copies of these reference genomes were removed from RefSeq b+f+h if present (See Supplementary Table 1 and 2 for detailed list of removals). The original read sets for ZYMO and ATCC samples can be found under SRA accessions ERR3152364 and SRR9847864 respectively.

#### Two Single-Organism Samples

Two pure samples, one containing *S. enterica* serovar Bareilly CFSAN000189, and the other containing *E. coli* O157:H7, were used to measure strain-level identification performance. Nanopore sequence data was available for both the *S. enterica* and *E. coli* samples under SRA accessions SRR9603470 and SRR9603471 respectively^32^. Each sample was sequenced by the MinION device, and the resultant read set was downsampled to 1Gb of sequence data for characterisation as above. For the *S. enterica* sample, all complete *S. enterica* genomes (875) available on RefSeq were downloaded and used as a reference database. For *E. coli,* the RefSeq b+f+h database was used.

### NanoMAP Program

#### Usage

Our implementation of the method, NanoMAP, is intended to be easy to use. The only requirements are related to the reference database against which the sample reads are characterised - each genome currently needs to be stored as a separate file, each header sequence must contain a human readable strain name as these appear in NanoMAP’s characterisation output, and the database must be non-redundant. A database with two or more copies of the reference genome for a sample strain will result in a scenario where no reads can be uniquely mapped, and all alignments have a MAPQ score of zero. A recommendation is to download genomes from an online database such as RefSeq, as this will satisfy all requirements (assume RefSeq is non-redundant).

#### Read classification after initial alignment

Filtering is performed prior to classification in order to remove low quality reads from consideration. Aside from removing short (< 1000 bp) reads, alignments with pid < 10%, or collinearity < 80% are removed. Here we define collinearity as the difference between the length of the query and that of the reference participating in the alignment, scaled by the length of the query participating in the alignment. This aims to remove alignments with large gaps.

After filtering, reads are classified. The classification for a read is a set of plausible strains the read may have originated from. For a single read, all alignments are gathered, then the following steps are performed to classify the read. The maximum alignment block (the alignment length between read and reference) is found, and any alignments with block < 50% of the maximum are removed. In a similar approach, the alignment with the highest number of base matches among those remaining is found, and alignments with base matches < 90% of this value are removed. Each remaining alignment has a high number of base matches and a large region of alignment for the particular read. The read is subsequently classified to the set of strains whose reference genomes are participating in these good quality alignments that remain.

#### Grouping Similar Strains

For each sample strain, a group of plausible candidate organisms is shortlisted. To facilitate this, the rate of classification co-occurrence for strains is created and consulted. A pairwise matrix is generated, which stores the frequency each strain appears in the same read classification as every other strain. Strains which are similar will possess a high classification co-occurrence rate, while those which are dissimilar will rarely appear together in a read classification and have low rate.

An abundance measure referred to as ‘naive abundance’ is used to sort strains by importance for the subsequent grouping process. For each read, each organism appearing in the read classification is awarded *n* bases, where *n* represents the length of the read divided by the number of organisms the read has been classified to. After this is calculated for each strain, the grouping process begins.

Strain grouping is performed using the pairwise classification co-occurrence matrix and the ordered list of strains. This is an iterative process, and continues till the ordered list has no remaining members. At each iteration, the strain with highest naive abundance is removed from the list and a new strain group is created for this organism. All remaining strains in the list are then collected into this group, where all strains with greater than 50% co-occurrence with the removed strain are added. The process then continues, where each time the organism with highest naive abundance is removed, then a group is formed around this strain.

#### Halving Candidates per Strain Group

After groups of candidate strains are formed, these groups usually need to be refined to a smaller set of organisms to ensure the MAPQ scores are useful. This is an iterative process, consisting of alignment, processing, then either halving the number of remaining strains or picking the present organisms. For a given strain group, an iteration begins by aligning the group’s read bin to a metagenome composed from reference genomes of the remaining strain candidates. The minimap2 alignment parameters are −c, −p 0.1, −N 10, −K 100M, −I 1000G, and read technology settings are applied with −x. After alignment, the output .paf file is processed in a similar manner to read classification after initial full database alignment, with some alterations. The classifications for a read are those with equal highest MAPQ score (highest MAPQ is often zero), and base matches equal to 99.9% of the highest base matches in alignments of the read. Naive abundance is calculated for each strain using the same method as previously mentioned. The number of MAPQ=60, MAPQ=10, and MAPQ=2 alignments for each strain are tallied for use when halving candidates or final strain identification.

Once the alignment file has been processed, the group of candidate strains is either halved in size, or final strain identification is performed. If the group has less than 5 strains remaining, or a single strain has MAPQ=60 alignment tally 100x greater than all others, the halving process is skipped and final identification is performed. If not, the group is halved using the following process. Strains with naive abundance of less than 0.05% of total sample abundance (5Mb of sequence data for 1Gb sample) are automatically removed. The new list of candidate strains is then created using MAPQ information. The two strains with highest MAPQ=60 alignment count are added to this new list if their MAPQ=60 alignment count is greater than 2. The same process is then repeated with MAPQ=10 alignments. If the size of the new candidate list is still less than half of the original value, strains are added in order of decreasing MAPQ=2 alignment count until the list is filled to the correct size. This forms the halved set of candidate strains for the group. A new metagenome is constructed from their reference genomes for alignment in the next iteration.

#### Picking Present Strains

Once the set of candidate strains is less than five, or a single strain is clearly evident, strains are selected as present in the sample given MAPQ information. If a single strain has a MAPQ=60 alignment count which is 10x greater than all others it is selected as the sole organism present for the group. Otherwise, any strain with a single MAPQ=60 alignment is selected as present. The strains identified in this manner appear in the final characterisation output of NanoMAP.

#### Strain Group Abundance Estimation

As summarized above, the probability model underlying our estimation procedure supposes that each read has an unknown probability *p*_*i*_ of being from strain *i* = *1,2,*…,*s*, where *s* is the number of strains in the strain group under consideration, and, given that a read is from strain *i*, it has probability *t*_*ij*_ of receiving classification *j,* where *j* =*1,2,* …*,2*^*s*^ −*1* indexes the possibly classifications of a read.

We first estimate the (transition) probabilities *{t_ij_}* by simulation. A ‘read set’ is created by extracting fragments of length 3000bp (constant stride) from each source genome. Fragments are marked with an identifier to indicate the genome from which the fragment was sampled. This read set is then aligned to a reference database consisting of all *s* source genomes, and the number of reads *m*_*ij*_ from source *i* receiving classification *j* is recorded. An estimated transition matrix is then constructed from this information: *t*_*ij*_ = *m*_*ij*_ / *Σ*_*k*_*m*_*ik*_, the proportion of reads from source genome *i* receiving classification *j*.

In practice, to obtain the classification frequencies for our read set, a reference database is built containing the genomes of just the identified strains (source genomes). Reads are aligned to this database and classified as described previously, where the classification is defined to be the set of strains participating in equal best alignments of the read. The frequencies *{n_j_}* observed in each classification are recorded and these are the data on which our estimates of proportions for the strain group will be based.

With the combination of the estimated transition matrix *{t_ij_}* and the observed classification counts *{n_j_}* for the read set, an EM-algorithm for estimating the probabilities *{p_i_}* can be readily derived (full details omitted). If after *c* iterations, we have estimates 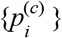, these are updated as follows:

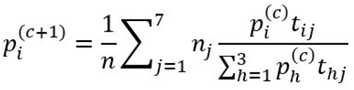

where *n* = *Σn*_*j*_ ·Natural starting values are 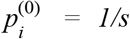, *i*=*1,*…*,s*. The EM iterations guarantee convergence as c → ∞ of the estimates 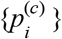 to stable values which give a *local maximum* of the observed data log-likelihood *LL*_*o*_, where

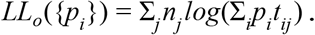

where here we assume that the matrix *{t_ij_}* has been estimated without error. If needed, different stable sets of estimates can be compared by calculating their respective *LL*_*0*_ values, and the set with the largest observed data log-likelihood retained. We have found that iterations starting from the values 1/s stabilized quickly at reasonable estimates of the *{p_i_}*, which we believe are the maximum likelihood estimates. So far we have not found evidence that multiple starting values need to be considered, though doubtless examples will be found in practice where this is the case.

#### Output

NanoMAP returns a strain-level characterisation against the reference genomes in an input genome folder. NanoMAP produces a simple output which lists the names, identifiers, and the sample DNA abundance of identified strains. The names and identifiers are those which appear in header lines of the input reference genomes. Most modern computers are suitable for use, with 8Gb of RAM being recommended. The most resource demanding part of NanoMAP is minimap2 alignment which can be configured for low system requirements to suit a given machine.

## Code Availability

NanoMAP and all associated code are available on GitHub (https://github.com/GraceAHall/NanoMAP).

## Acknowledgements

This work has been supported by an Alan W Harris scholarship provided by the Walter and Eliza Hall Institute of Medical Research. Special thanks to Leonard Harrison for the use of lab space, Wei Shi and Yang Liao for discussions on their tool sublong, and to Matthew Ritchie and Quentin Gouil for consulting and assisting during the project.

