## Supplementary Information for "Strain-level sample characterisation using long reads and MAPQ scores"

### Estimating Strain Discrimination Sensitivity

Here we provide an estimate of the sensitivity of strain discrimination using the method developed in this paper, and using indicative parameters for sequence data that would be obtained by the current state-of-the-art ONT flongle device. We consider a pair of strains, $g_1$ and $g_2$. Structural variants are those contiguous sequences of 50 or more bases that are not common to both genomes, or are reversed or translocated with respect to each other. The derivation here is similar to that of Lander and Waterman[33] that was developed for a rather different purpose and context. We seek an expression for the probability that at least one SV boundary is overlapped adequately by a read that is well-aligned to exactly one of a pair of genomes. Since we are focused on the lower limits of sensitivity, we consider only pairs of highly similar genomes that differ in having a small number (e.g. <10) of structural variants relative to each other.

Denote by G denote the genome length of the alternate in bases; R, the read length, assumed fixed here, but - in reality - typically unimodally distributed with positive skewness; and V the length of the structural variant. Let the minimum required overlap of the read into both the structural variant region and the common region of the read with respect to the alternate genome be of length C. We assume $C < V < R$.

Suppose we are interested in the sensitivity to identifying the strain $g_1$.

Let N be the number of sequenced bases of DNA fragments derived from the microbiome sample. Let s be the fraction of $g_1$ present in the sequenced microbiome sample, where that fraction is in terms of bacterial cells (and hence copies of the genome).

Define a *valid* read as one which lies wholly within the genome, and *overlap* of a read with an SV to mean the read has at least C bases within the SV and also within the adjacent region of the two genomes. While we assume here that all SVs involve additions into genome $g_1$, the logic is sound for deletions, which are SVs of length 0 in $g_1$, but length $V>0$ in $g_2$.

Consider the case of there being a single SV. Let the 5' end of a particular read be at location x on the genome, and the 5' end of the SV at y. Then x lies in [1, G-R], and y lies in [1, G-V]. There are two very minor end constraints – for a read to overlap the 5' end of an SV $y \geq C$, and to overlap the 3' end $y < G-C$. The latter constraint is redundant given that $y \leq G-V$. We will ignore the other end constraint. Provided $y \geq R-C$ and $y \leq G-V$ any read that lies in [y-R+C, y+V-C] will overlap the SV. This corresponds to a length of R+V-2C along the genome that would give a read overlapping the SV. Assume that SVs are independently and uniformly located along the genome, and that their 5' ends lie within [1, …, G-V]. Then the probability that a randomly located valid read does not overlap this SV is

$$\text{Prob(no overlap \& valid)} = \text{Prob(no overlap | valid) Prob(valid)} = \frac{G-(R+V-2C)}{G-R}\frac{G-R}{G} = 1 - \frac{R+V-2C}{G}$$

Since $C \ll R$ we will ignore it in the remainder of this development.

Now consider the case of multiple SVs. A further approximation is made – namely that the SVs are mutually independent and uniformly distributed along the genome (except near the genome ends), and constrained to not be within a distance of each other that allows a read to overlap more than one SV. Following[34] it can be shown that this condition would be met in more than 99.5% of cases for the R and V values considered here. With this condition overlap of each of the SVs by a read are mutually exclusive events. Hence the probability of overlapping one SV given there are n SVs present is $\frac{n(R+V)}{G}$ ,

and the probability of no overlap of any of the SVs by a single read is 1- $\frac{n(R+V)}{G}$

The total number of reads is $N/R$ , and the expected number of reads from the strain is sN/R, where s is the fraction of reads from strain $g_1$. Assuming that these reads are uniformly distributed along the genome, the probability of at least one read from the microbiome sample crossing at least one boundary that discriminates this particular strain from a genome that differs from it by n structural variants is

$$p_{1hit} = 1 - \left(1 - \frac{n(R-V)}{G}\right)^{sN/R}$$

Figure S.1 illustrates the variation of the probability of detecting at least one differentiating boundary between the strain and the reference as a function of various parameters. Here we have considered the case G = 3Mb, R = 10Kb, C= 100, V = 2000, for a range of values of N, s, and n. From this we see that, under the assumptions of this model, with 1 Gb of sequence data a strain present at 0.25% and differing from another strain by a single structural variant would be expected to give clear MAPQ evidence of its presence in more than 50% of cases. Noting that strains differing by as few as 5 SVs are not common, it can be seen that - with only 200Mb of sequence data - a comparable performance would be achieved for an organism only 5 SVs different from another.

In reality, read coverage of any genome will not be uniform. Consequently, there will be regions where coverage will be less than the mean. The plots of Figure S.1 allow one to estimate the effect of non-uniformity-induced reduced coverage on the limits of strain sensitivity. This would correspond to lower levels of coverage – for instance, to the coverage being halved, and hence moving from one curve (corresponding, for example, to 1Gb of data) to the one corresponding to 0.5Gb of data. The likely dominant factor that our simple model fails to consider is the non-uniformity of coverage rather than other approximations – such as genome end effects – that have been explicitly stated.

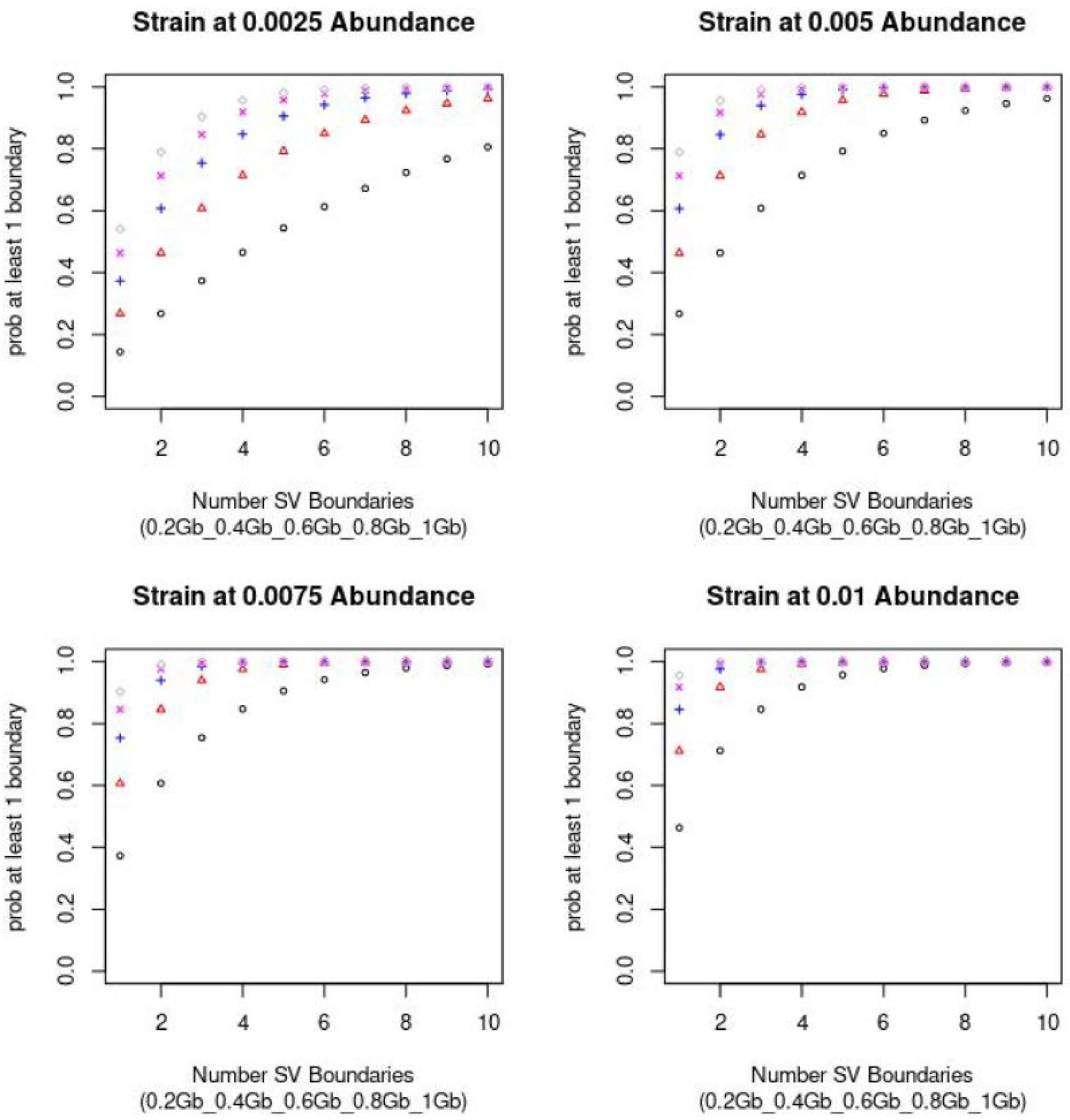


Figure S.1. Probability of finding at least one structural variant boundary when comparing reads against a reference genome that has from 1 to 10 structural variants relative to the organism from which the reads actually came. Each sub-figure gives probability variation with number of SVs assuming sequencing providing 5 levels of base counts (0.2Gb to 1.0Gb), with the sample strain present at a relative abundance of either 0.0025, 0.005, 0075, or 0.01. It is assumed that reads are of length 10000 bases, that the required overlap of the read into the variant region and the common region is at least 100, and that the structural variant is 2000 bases long.

### Database Makeup for Zymo and Atcc Sample Characterisation

As mentioned in the main text, the RefSeq b+f+h database consists of all available complete RefSeq bacterial and fungal genome assemblies with the addition of hg38. The organisms present in the ZYMO and ATCC samples were added to RefSeq b+f+h for characterisation of these products. This process introduced redundant copies of the reference genome for sample organisms in some cases. Supplementary tables 1 and 2 outline the organisms that were removed for ATCC and ZYMO characterisation.

| Sample Organism | Reference Genome added to database | Redundant Copies removed from database |
|---|---|---|
| *B. fragilis* ATCC 25285 | ATCC reference genome | GCF_000025985.1, GCF_005706655.1 |
| *B. vulgatus* ATCC 8482 | ATCC reference genome | GCF_000012825.1 |
| *B. adolescentis* ATCC 15703 | ATCC reference genome | GCF_000010425.1 |
| *C. difficile* ATCC 9689 | ATCC reference genome | GCF_001077535.1, GCF_002073735.2 |
| *E. faecalis* ATCC 700802 | ATCC reference genome | GCF_000007785.1 |
| *L. plantarum* ATCC BAA-793 | ATCC reference genome | GCF_000203855.3 |
| *E. cloacae* ATCC 13047 | ATCC reference genome | GCF_000025565.1, GCF_013376815.1 |
| *E. coli* ATCC 700926 | ATCC reference genome | - |
| *H. pylori* ATCC 700392 | ATCC reference genome | GCF_000008525.1 |
| *S. enterica* ATCC 9150 | ATCC reference genome | GCF_000011885.1 |
| *Y. enterocolitica* ATCC 27729 | ATCC reference genome | GCF_000834195.1 |
| *F. nucleatum* ATCC 25586 | ATCC reference genome | GCF_003019295.1 |

Table S.1. Reference genomes for ATCC sample strains added to RefSeq b+f+h, and redundant genome assemblies removed. For each sample strain, its ATCC reference genome was downloaded from the ATCC genome portal (https://www.atcc.org/Landing_Pages/Genome_Portal.aspx). Many of these ATCC genome assemblies are already present in RefSeq, requiring their RefSeq submission to be removed from the database prior to NanoMAP characterisation.

| Sample Organism | Reference Genome in Database | Removed Redundant Copies |
|---|---|---|
| *B. subtilis* ZymoBIOMICS | ZymoBIOMICS reference genome | GCF_006364795.1 |
| *E. faecalis* ZymoBIOMICS | ZymoBIOMICS reference genome | GCF_006364815.1 |
| *E. coli* ZymoBIOMICS | ZymoBIOMICS reference genome | GCF_006364035.1 |
| *L. fermentum* ZymoBIOMICS | ZymoBIOMICS reference genome | GCF_006094475.1, GCF_006364495.1 |
| *L. monocytogenes* ZymoBIOMICS | ZymoBIOMICS reference genome | - |
| *P. aeruginosa* ZymoBIOMICS | ZymoBIOMICS reference genome | - |
| *S. enterica* ZymoBIOMICS | ZymoBIOMICS reference genome | - |
| *S. aureus* ZymoBIOMICS | ZymoBIOMICS reference genome | - |

Table S.2. Reference genomes for ZymoBIOMICS sample strains added to RefSeq b+f+h, and redundant genome assemblies removed. All reference genomes for Zymobiomics strains were downloaded from https://s3.amazonaws.com/zymo-files/BioPool/ZymoBIOMICS.STD.refseq.v2.zip. Half the ZymoBIOMICS strains had FDAARGOS assembly submissions in RefSeq, requiring them to be removed prior to NanoMAP characterisation.
